# Molecular Contrastive Learning with Graph Attention Network (MoCL-GAT) for Enhanced Molecular Representation

**DOI:** 10.1101/2025.09.17.676724

**Authors:** Alperen Dalkıran, Ahmet S. Rifaioğlu, Rengul Cetin-Atalay, Aybar C. Acar, Tunca Doğan, M. Volkan Atalay

**Affiliations:** Department of Computer Engineering, Middle East Technical University, 06800, Ankara, Türkiye; Department of Computer Engineering, Adana Alparslan Türkeş Science and Technology University, 01250, Adana, Türkiye; Heidelberg University, Faculty of Medicine, and Heidelberg University Hospital, Institute for Computational Biomedicine, Heidelberg, Germany; Department of Medicine, Section of Pulmonary and Critical Care Medicine, The University of Chicago, Chicago, IL 60637, U.S.A.; Cancer Systems Biology Laboratory (KanSiL), Graduate School of Informatics, Middle East Technical University, 06800, Ankara, Türkiye; Biological Data Science Lab, Department of Computer Engineering, Hacettepe University, 06800, Ankara, Türkiye; Department of Bioinformatics, Graduate School of Health Sciences, Hacettepe University, 06800, Ankara, Türkiye; Deptartment of Health Informatics, Institute of Informatics, Hacettepe University, 06800, Ankara, Türkiye; Department of Information Systems and Supply Chain Management, Loyola University Chicago, Chicago, IL 60611, U.S.A.

## Abstract

Learning the representation of molecules is crucial for drug discovery but is often hindered by the scarcity of labeled experimental data, which limits the performance of supervised machine learning models. While self-supervised learning (SSL) offers a solution by leveraging vast unlabeled chemical databases, many existing methods focus on learning from either local structural information or global molecular properties, but not both simultaneously. We introduce MoCL-GAT, a novel contrastive and transfer learning-based SSL framework that addresses this gap by simultaneously learning from two complementary objectives. It combines a local contrastive task on molecular subgraphs to capture fine-grained chemical environments with a global predictive task to learn holistic molecular descriptors. This dual-objective approach, powered by a Graph Attention Network, is designed to create more robust, versatile, and transferable molecular representations. Pre-trained on 1.9 million compounds, MoCL-GAT was fine-tuned on diverse benchmarks. It achieved state-of-the-art performance on molecular property prediction tasks, with an AUROC of 0.928 on BBBP and 0.749 on SIDER, and top-ranking RMSEs of 0.570 for ESOL and 1.818 for FreeSolv. Critically, fine-tuned models consistently and significantly outperformed models trained from scratch, confirming the value of pre-training. These results validate that MoCL-GAT’s dual-objective approach learns highly effective and transferable representations, enabling more accurate and data-efficient predictions for key cheminformatics challenges.

## 1 Introduction

Computational approaches are increasingly pivotal in modern drug discovery, offering powerful tools to accelerate the identification of novel therapeutics by predicting molecular properties and drug-target interactions (DTIs) (Schneider, 2018; Rifaioglu *et al*., 2019, 2020). The effectiveness of these in silico methods depends on developing informative molecular representations that capture essential chemical and structural features. Graph Neural Networks (GNNs), and particularly Graph Attention Networks (GATs) (Veličković *et al*., 2018), have become a leading architecture for this task, as their message-passing paradigm naturally aligns with the graph structure of molecules, enabling the learning of intricate local atomic patterns.

Despite their power, the standard supervised training of GNNs is often limited by the high cost and low availability of large-scale labeled experimental datasets. This data scarcity is a critical bottleneck, especially for novel chemical scaffolds or understudied biological targets, limiting the ability of supervised models to generalize across the vast chemical space (Dalkıran *et al*., 2023). Self-supervised learning (SSL) provides a powerful solution to this challenge by learning transferable representations from massive unlabeled databases (Liu *et al*., 2021). By designing pretext tasks where the supervision signal is derived from the data itself, SSL models can learn fundamental chemical principles that significantly boost performance when later fine-tuned on specific, smaller labeled datasets (Rong *et al*., 2020).

Recent SSL approaches for molecular representation have largely followed several key trends. One major direction involves adapting Transformers-based language models, such as BERT, to chemical notations e.g., SMILES or SELFIES (Liu *et al*., 2019; Fabian *et al*., 2020; Ahmad *et al*., 2022; Yüksel *et al*., 2023). Another prominent direction focuses on graph-based methods, utilizing contrastive or generative objectives on 2D molecular graphs, such as predicting masked atomic attributes or contrasting different augmented views of a molecule (Hu *et al*., 2020; Wang *et al*., 2022). More recently, methods have incorporated 3D geometric information to learn richer, conformation-aware embeddings (Liu *et al*., 2022; Fang *et al*., 2022; Chen *et al*., 2025) or leverage existing domain knowledge like chemical descriptors to guide the pre-training process (Schütt *et al*., 2017; Lu *et al*., 2019; Li *et al*., 2022). While each of these strategies has proven successful, they often emphasize learning signals from a single modality or scale. (A detailed review of related works is provided in the Supplementary Material, Section S1).

Despite progress, a major challenge remains in developing a single framework. This framework must be capable of learning from both the detailed local chemical environments that govern specific interactions and the broader global properties that influence a molecule’s overall behavior. Models that focus solely on local contrastive tasks may fail to capture essential global features, such as solubility or membrane permeability. Similarly, models that only predict global properties may overlook critical subgraph motifs necessary for binding affinity. Therefore, a key unsolved problem is developing a self-supervised approach that explicitly integrates these multi-scale representations in order to produce more comprehensive and chemically varied embeddings.

In this context, we introduce Molecular Contrastive Learning with Graph Attention Network (MoCL-GAT), a novel self-supervised learning framework specifically designed to generate comprehensive and robust molecular representations using GATs. MoCL-GAT addresses the need to capture information at multiple scales by integrating learning objectives. These objectives are derived both from local atomic environments as well as from global molecular characteristics within a unified dual self-supervised strategy. We hypothesize that this dual approach is crucial for effective molecular modeling: while local structure dictates chemical reactivity and specific binding interactions, global properties influence aspects such as solubility, membrane permeability, and overall drug-likeness. MoCL-GAT operationalizes this concept through two complementary components: (1) applied to K-hop subgraphs augmented via attribute masking, forcing the model to learn representations invariant to local perturbations while preserving essential neighborhood structures, and (2) a global prediction task where the model learns to predict a curated set of established RDKit (Landrum, 2006) molecular descriptors, encouraging the embeddings to summarize overall molecular features. The GAT architecture serves as the backbone encoder, processing the graph information to support both learning objectives. By combining these strategies, MoCL-GAT aims to produce molecular embeddings that are both chemically meaningful and broadly transferable across downstream tasks.

The primary contributions of this work are threefold:

1. **A Novel Dual SSL Framework:** We introduce and implement MoCL-GAT, a self-supervised learning strategy that explicitly integrates local structural learning via contrastive methods with global property learning via molecular descriptor prediction-specifically tailored for molecular graph data.
2. **An Integrated Multi-Scale Learning Approach:** We detail the effective integration of key components within the MoCL-GAT framework, including K-hop subgraph sampling, attribute masking, Information Noise-Contrastive Estimation (InfoNCE) (Oord *et al*., 2019), global descriptor regression, and the use of GAT architecture as the backbone encoder within the MoCL-GAT framework.
3. **State-of-the-Art Performance:** Through extensive empirical evaluation, we demonstrate that pre-training with MoCL-GAT yields highly transferable molecular representations. Upon fine-tuning, these embeddings lead to significant performance gains and achieve state-of-the-art or highly competitive results across a wide array of standard molecular property prediction and DTI-related benchmarks.

This work highlights the effectiveness of integrating multi-scale self-supervised signals for learning generalizable molecular representations. Our work demonstrates that pre-training on large unlabeled datasets with a dual-objective strategy is an effective approach for developing accurate predictive models, particularly when labeled data for specific downstream tasks is limited. The subsequent sections elaborate on the details of the methodology (Section 2), present comprehensive experimental results and comparative analyses (Section 3), discuss the findings and their implications (Section 4), and conclude by outlining potential directions for future research (Section 5).

## 2 Methods

### 2.1 Overview of the MoCL-GAT Framework

The Molecular Contrastive Learning with Graph Attention Network (MoCL-GAT) framework adopts a two-stage learning paradigm, comprising self-supervised pre-training followed by supervised fine-tuning. The primary objective of the pretraining stage, illustrated in **Figure 1a**, is to learn rich, transferable molecular representations from large-scale unlabeled datasets by optimizing a novel dual self-supervised objective function. These representations are then adapted during the fine-tuning stage (**Figure 1b**) to specific downstream predictive tasks using corresponding labeled datasets. The primary objective of the pre-training stage is to learn rich, transferable molecular representations from large-scale unlabeled datasets by optimizing a novel dual self-supervised objective function. These representations are then adapted during the fine-tuning stage to specific downstream predictive tasks using corresponding labeled datasets. The core of MoCL-GAT is a Graph Attention Network (GAT) (Veličković *et al*., 2018) encoder. GATs leverage attention mechanisms to dynamically assign importance to neighboring atoms, enabling the model to focus on the most relevant local context during representation learning.

**Figure 1.**
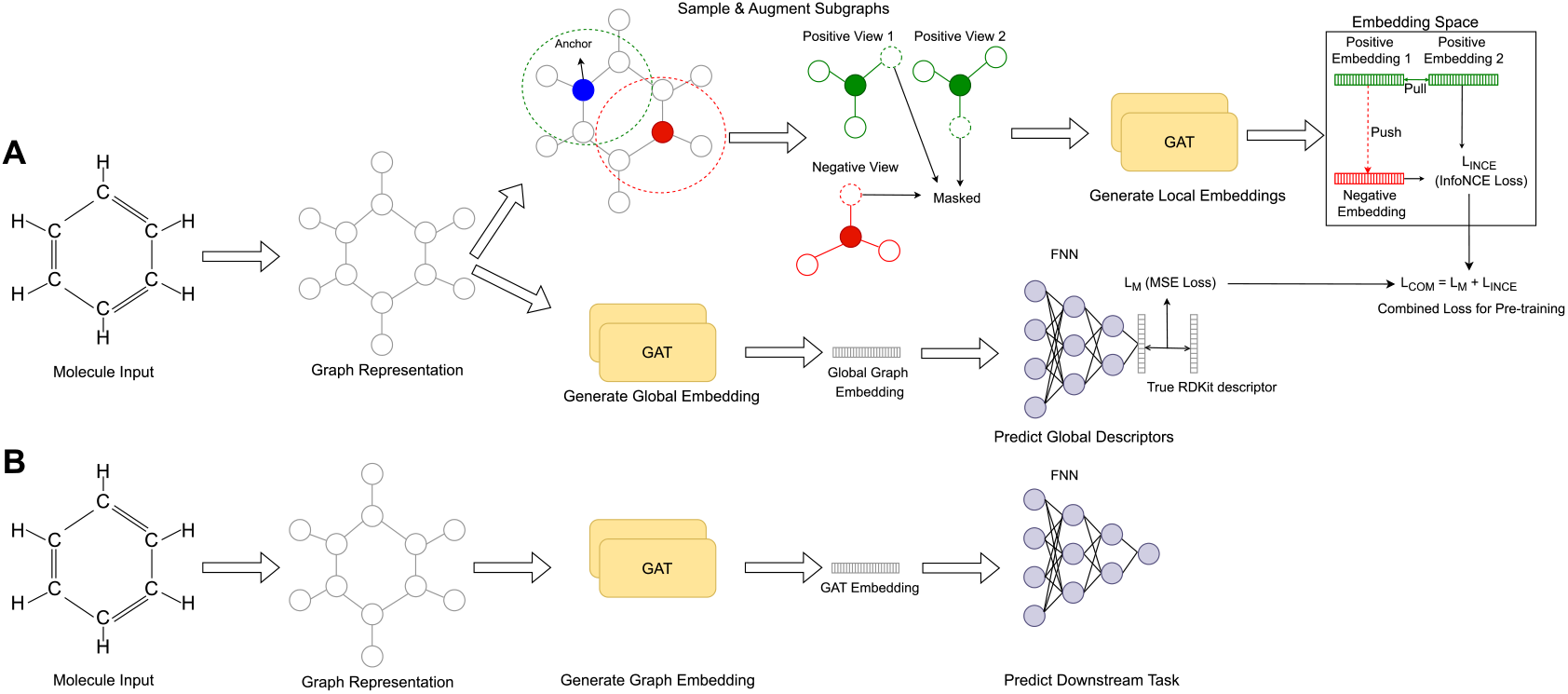
Overview of the MoCL-GAT framework. **A)** The self-supervised pre-training phase operates on two parallel objectives. In the local pathway (top), an anchor node (blue) and a negative node (red) are sampled from the graph. Augmented views are generated from their respective subgraphs, creating a positive pair and a negative view. These are processed by a GAT to generate local embeddings. A contrastive loss (*L*_*INCE*_) is then applied within the embedding space to attract positive pairs and push them from negative ones. In the global pathway (bottom), the entire graph is processed by a GAT to produce a global graph embedding, which is used by an FFN to predict a vector of RDKit descriptors. The Mean Squared Error loss (*L*_*M*_) is calculated against the true descriptors. Both losses are combined (*L*_*COM*_) to train the model. **B)** In the supervised fine-tuning phase, the pre-trained GAT encoder generates embeddings from input molecules, which are fed into a task-specific FFN for downstream predictions.

### 2.2 Molecular Graph Representation

Input molecules, initially provided as Simplified Molecular Input Line Entry System (SMILES) strings, were converted into graph-based representations suitable for processing by GNNs. This conversion was performed using the RDKit cheminformatics library. Each molecule is represented as a graph *G = (V, E)*, where the node set *V* corresponds to atoms and the edge set *E* corresponds to chemical bonds within the molecule.

### 2.3 Feature Extraction

To initialize the molecular graph with chemically meaningful information, features were extracted at multiple levels: atom (node), bond (edge), and molecule (global).

#### 2.3.1 Node (Atom) Features

Each node *v* ∈ *V* was initialized with a feature vector summarizing its atomic properties. These features consist of: atomic number, chirality tag (R/S configuration), degree (connectivity), formal Charge, count of explicit hydrogen atoms attached, count of radical electrons, hybridization state (e.g., sp, sp^2^, sp^3^), aromaticity status (boolean), and ring membership (boolean indicator of participation in any ring structure).

#### 2.3.2 Edge (Bond) Features

Each edge *e* ∈ *E*, representing a chemical bond, is decorated with features describing its characteristics. These features include: bond type (single, double, triple, or aromatic), bond stereochemistry (e.g., cis/trans, E/Z configuration), and conjugation status (boolean indicator of participation in a conjugated system).

#### 2.3.3 Global (Molecule-level) Descriptors

A curated set of molecule-level descriptors were computed using RDKit to serve as targets for the global self-supervised task. This descriptor set was acquired from the GROVER framework (Rong *et al*., 2020) and further refined through empirical analysis. Initially, a set composed of 85 descriptors was considered; however, 36 descriptors with values consistently below 0.00001 were excluded due to their negligible contribution to representation learning. To enhance the expressiveness of the global molecular embeddings, an additional 29 descriptors—drawn from diverse chemical property categories, including the Synthetic Accessibility (SA) Score (Ertl and Schuffenhauer, 2009) were incorporated. This refinement resulted in a final set of 78 informative molecular descriptors, designed to capture a wide range of physicochemical, topological, and structural properties relevant to drug discovery. The full list of descriptors can be found in Table S1.

### 2.4 Self-Supervised Pre-training Strategy

The pre-training phase is designed to learn rich molecular representations by simultaneously optimizing objectives related to both local graph structure and global molecular properties, using only unlabeled data.

#### Local Perspective (Contrastive Learning)

This component focuses on capturing fine-grained local chemical environments through a contrastive learning approach applied to K-hop subgraphs. For a randomly selected anchor node *v*, its 2-hop subgraph is extracted. Two distinct augmented views of this subgraph are generated by randomly masking node and edge attributes, forming a positive pair. Negative samples are generated by selecting subgraphs centered on nodes *u* outside the 2-hop neighborhood of *v*. All augmented views are processed by the GAT encoder to produce embeddings.

The objective is to maximize the agreement between embeddings of positive pairs while minimizing it for negative pairs, which is formalized using a variant of the Information Noise-Contrastive Estimation (InfoNCE) loss. Given an anchor subgraph embedding *h*_*i*_, its positive pair embedding *h*_*j*_, and a set of *N* negative embeddings {*h*_*k*_}_k=1..N_, the contrastive loss is defined as:

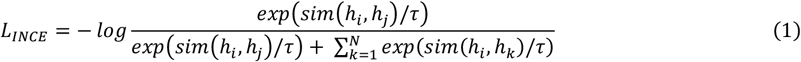

where *sim(u, v)* is the cosine similarity between vectors *u* and *v*, and τ is a temperature hyperparameter that scales the distribution of similarities, which we set to 0.5. Minimizing this loss encourages the model to pull representations of positive pairs closer together in the embedding space while pushing them apart from negative samples.

#### Global Perspective (Descriptor Prediction)

In parallel, the model learns to capture global molecular characteristics. Node embeddings generated by the GAT encoder for the entire molecular graph are aggregated into a single graph-level representation *h*_*G*_ using global mean pooling. This graph embedding is then fed into the FFN prediction head to predict the vector of 78 pre-computed RDKit descriptors (Section 2.3.3).

This predictive task is optimized by minimizing the Mean Squared Error (MSE) loss, denoted as *L*_*M*_, between the predicted and true descriptor values. Given a predicted descriptor vector *ŷ* and the true descriptor vector *y*, both of dimension *D* (where *D*=78), the loss is defined as:

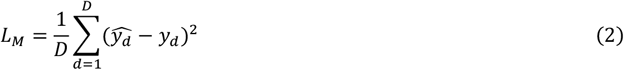

This objective function encourages the GAT encoder to produce graph embeddings that retain holistic information about the molecule’s physicochemical and structural properties.

#### Combined Pre-training Objective

The final objective function for the self-supervised pre-training phase is a direct combination of the local contrastive loss and the global predictive loss. The model is trained end-to-end by minimizing this combined loss, *L*_*COM*_, which equally weights both tasks:

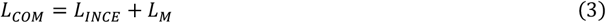

This dual objective ensures that the learned representations are simultaneously enriched with knowledge of both local chemical environments and global molecular properties, leading to a more comprehensive and versatile foundation for downstream tasks.

### 2.5 Pre-training Dataset

Our study utilizes data from the ChEMBL (Gaulton *et al*., 2012) database version 33, a comprehensive repository of bio-active molecules with drug-like properties. After preprocessing, 1,915,183 compounds with valid SMILES representations were retained from an initial collection of 1,920,643 compounds. This large-scale dataset provides a robust foundation for developing and training our molecular contrastive learning approach.

### 2.6 Supervised Fine-tuning

Following the self-supervised pre-training phase, the learned parameters of the GAT encoder served as a starting point for supervised fine-tuning on specific downstream tasks. The benchmark datasets used for evaluation, corresponding to the results presented in **Error! Reference source not found**., included classification tasks binding properties against the human beta-secretase 1 protein (BACE), the blood–brain barrier penetration (BBBP), the ability to inhibit HIV replication (HIV), the “Toxicology in the 21st Century” (Tox21), the Side Effect Resource (SIDER), and regression tasks aqueous solubility (ESOL), The Free Solvation Database (FreeSolv), Lipophilicity, the binding affinity prediction (PDBbind), primarily sourced from the MoleculeNet collection (Wu *et al*., 2018). For MoleculeNet datasets, standard scaffold-based splitting methodologies were used to ensure robust evaluation on novel chemical scaffolds. During fine-tuning, the entire MoCL-GAT model, including the pre-trained GAT encoder and the task-specific FFN prediction head, was trained end-to-end on the labeled data using task-appropriate loss functions (i.e., Binary Cross-Entropy for classification, MSE for regression).

### 2.7 Evaluation Metrics

Performance was assessed using standard metrics relevant to each task type. For classification benchmarks, the Area Under the Receiver Operating Characteristic curve (AUROC) was reported. For regression benchmarks, Root Mean Squared Error (RMSE) was used as the metric.

### 2.8 Model Architecture

MoCL-GAT’s GAT encoder is composed of two GAT layers. We performed a comprehensive hyperparameter optimization using a grid search strategy to determine the optimal model configuration. The full search space for all tested parameters is detailed in Table S2. The optimization process identified the optimal configuration as follows: a learning rate of 0.0001, a dropout rate of 0.2, a batch size of 128, and 16 attention heads for the GAT encoder. The outputs from the multiple attention heads in the final GAT layer were averaged before subsequent processing.

The output of the final GAT layer was then passed into a two-layer feed-forward network (FFN) with hidden dimensions of 128 and 256. During pre-training, this FFN functioned as the prediction head for global RDKit descriptors. In the fine-tuning phase, a similarly structured FFN was employed as the task-specific prediction head, incorporating appropriate activation functions (e.g., ReLU) and output units (e.g., linear for regression, sigmoid for binary classification).

### 2.9 Implementation Details

The MoCL-GAT framework was implemented in Python, leveraging the PyTorch library (Paszke *et al*., 2019) for deep learning model construction. Graph-specific operations and GNN layers were implemented using the PyTorch Geometric (PyG) library (Fey and Lenssen, 2019). Cheminformatics tasks, including molecule parsing, feature extraction, and descriptor calculation, were handled using the RDKit library.

## 3 Results

To assess the effectiveness of the proposed MoCL-GAT framework, we conducted comprehensive experiments comparing its performance against state-of-the-art baseline methods on a diverse set of established molecular benchmark datasets. The evaluation encompassed both classification and regression tasks relevant to drug discovery, with a focus on assessing the quality and transferability of the molecular representations learned during the self-supervised pre-training phase.

**Table 1.** Comparative Performance Analysis of MoCL-GAT and Baseline Models via Scaffold Splitting on Standard Benchmark Datasets.

|  | AUROC (Higher is better) |  |  |  |  | RMSE (Lower is better) |  |  |  |
| --- | --- | --- | --- | --- | --- | --- | --- | --- | --- |
| Method | BACE | BBBP | HIV | Tox 21 | SIDER | ESOL | FreeSolv | Lipophilicity | PDBbind |
| SELFormer | 0.832 | 0.902 | 0.681 | 0.653 | 0.745 | 0.682 | 2.797 | 0.735 | 1.488 |
| ChemProp | 0.809 | 0.710 | 0.771 | 0.759 | 0.570 | 1.050 | 2.082 | 0.683 | **1.397** |
| MolBERT | 0.866 | 0.762 | 0.783 | - | - | - | - | - | - |
| ChemBERTa-2 | 0.799 | 0.728 | 0.622 | - | - | - | - | 0.986 | - |
| Hu et al. | 0.859 | 0.708 | 0.802 | 0.787 | 0.652 | 1.220 | 2.830 | 0.740 | - |
| MolCLR | **0.890** | 0.736 | **0.806** | 0.787 | 0.652 | 1.110 | 2.200 | 0.650 | - |
| GraphMVP-C | 0.812 | 0.724 | 0.770 | 0.744 | 0.639 | 1.029 | - | 0.681 | - |
| GEM | 0.856 | 0.724 | **0.806** | 0.781 | 0.672 | 0.798 | 1.877 | 0.660 | - |
| MGCN | 0.734 | 0.850 | 0.738 | 0.707 | 0.552 | 1.270 | 3.350 | 1.110 | - |
| GCN | 0.716 | 0.718 | 0.740 | 0.709 | 0.536 | 1.430 | 2.870 | 0.850 | - |
| GIN | 0.701 | 0.658 | 0.753 | 0.740 | 0.573 | 1.450 | 2.760 | 0.850 | - |
| SchNet | 0.766 | 0.848 | 0.702 | 0.772 | 0.539 | 1.050 | 3.220 | 0.910 | - |
| KPGT | 0.855 | 0.908 | - | **0.848** | 0.649 | 0.803 | 2.121 | **0.600** | - |
| MolGT | 0.845 | 0.737 | 0.793 | 0.758 | 0.654 | 0.839 | - | 0.788 | - |
| MoCL-GAT (ours) | 0.821 | **0.928** | 0.768 | 0.740 | **0.749** | **0.570** | **1.667** | 0.684 | 1.456 |

### 3.1 Performance on Benchmark Datasets

The primary quantitative evaluation involved fine-tuning the pre-trained MoCL-GAT model on various downstream tasks and comparing its performance against baseline models, including SELFormer, D-MPNN, MolCLR, GEM, and KPGT. **Error! Reference source not found**. summarizes the performance metrics – Area Under the Receiver Operating Characteristic curve (AUROC) for classification tasks and Root Mean Squared Error (RMSE) for regression tasks. Lower RMSE and higher AUROC values indicate better performance.

As shown in **Error! Reference source not found**., MoCL-GAT demonstrates strong performance across the evaluated benchmarks achieving state-of-the-art results on several key tasks. In classification, MoCL-GAT attains the highest AUROC scores on the BBBP (0.928) and SIDER (0.749) datasets, indicating superior capability in predicting blood-brain barrier penetration and compound side effects, respectively. While other methods such as MolCLR and KPGT exhibit strengths on tasks like BACE, HIV, and Tox21, MoCL-GAT remains highly competitive on these benchmarks.

In the regression domain, MoCL-GAT significantly outperforms baseline methods in predicting water solubility (ESOL, RMSE = 0.570) and hydration free energy (FreeSolv, RMSE = 1.667), achieving the lowest errors among all compared models. Its performance on Lipophilicity (RMSE = 0.684) and PDBbind (RMSE = 1.456) is also competitive, tough slightly surpassed by KPGT and D-MPNN, respectively, on these specific tasks. Overall, the results across this diverse set of benchmarks highlight the effectiveness of MoCL-GAT in learning generalizable molecular representations. These representations are applicable to a wide range of molecular modeling challenges, encompassing both pharmacokinetic properties (BBBP, ESOL, Lipophilicity) and specific bioactivities or molecular interactions (SIDER, HIV, BACE, Tox21, PDBbind).

### 3.2 Visualization of Learned Embeddings

To gain qualitative insights into the structure of the learned representations, we visualized the molecular embeddings using t-distributed Stochastic Neighbor Embedding (t-SNE) (Van der Maaten and Hinton, 2008). **Figure *2*** presents the t-SNE plots of molecule embeddings for two representative binary classification datasets BACE (top row) and BBBP (bottom row), at different stages of the learning process. To quantitatively evaluate the separability of these embeddings, we applied Agglomerative Hierarchical Clustering and calculated the resulting cluster purity scores.

**Figure 2.**
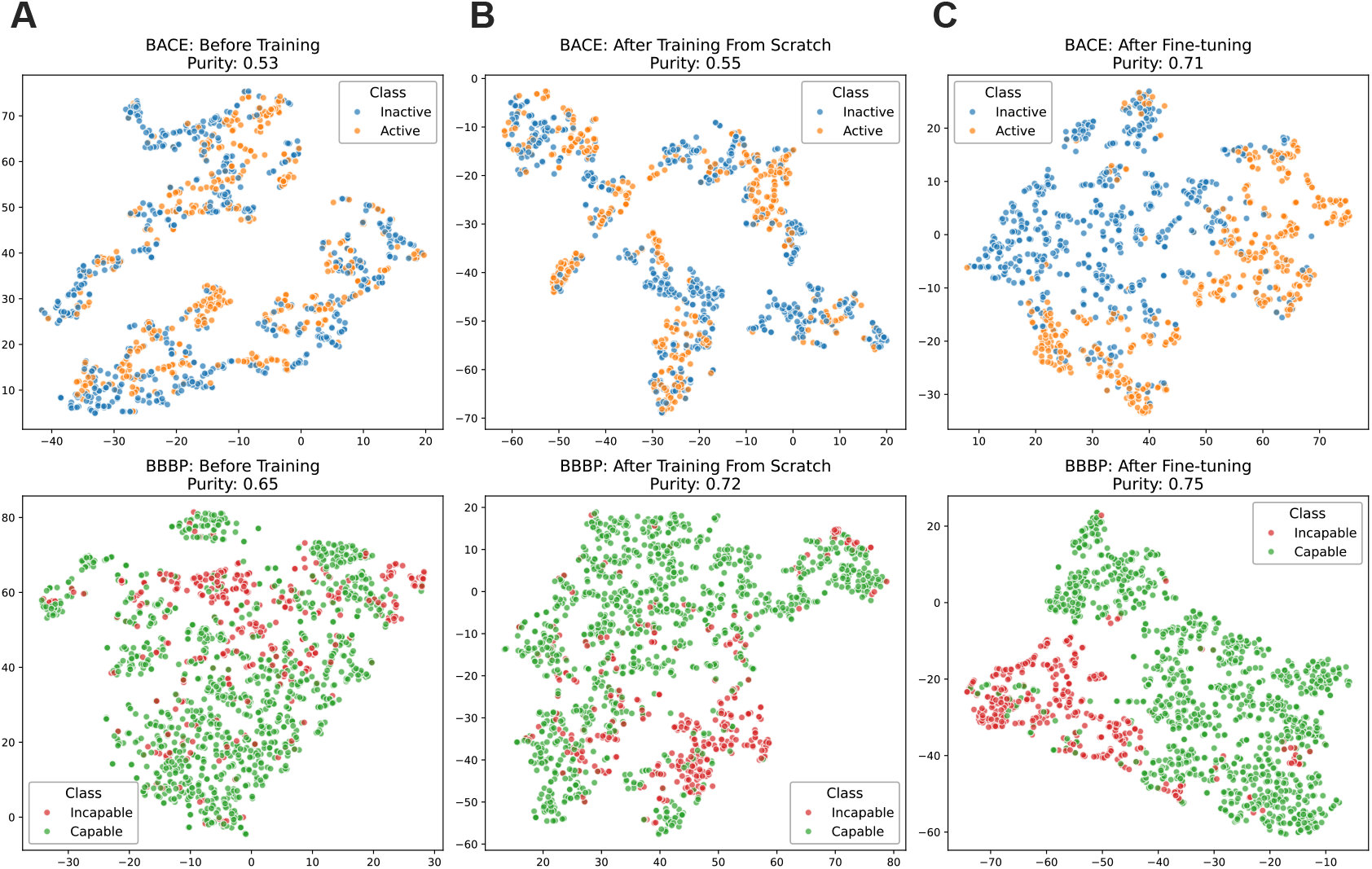
t-SNE visualization of molecular embeddings for the BACE (top row) and BBBP (bottom row) datasets at different learning stages. **A)** In the initial state, embeddings for both BACE (top-left) and BBBP (bottom-left) show no effective separation between classes. This is reflected in low purity scores of [0.53] and [0.65], respectively, indicating that active/inactive and capable/incapable molecules are highly intermixed. **B)** Following the MoCL-GAT training from scratch phase, the embeddings begin to show emergent structure based on general chemical features. This results in a moderate improvement in cluster purity for both BACE ([0.55]) and BBBP ([0.72]). **C)** After supervised fine-tuning on each specific task, the embeddings exhibit significant separability. Distinct clusters are formed for BACE inhibitors (orange) vs. non-inhibitors (blue) and for BBBP-capable (green) vs. incapable (red) compounds. This qualitative visual improvement is quantitatively confirmed by the highest purity scores of the process: [0.71] for BACE and [0.75] for BBBP.

A. Before Training: The leftmost panels show the t-SNE visualization of initial embeddings prior to any training. As expected, molecules from to the two classes are largely intermixed, indicating no inherent structure or separation in the initial representation. This lack of separation is quantitatively confirmed by low initial purity scores for both datasets (e.g., [0.53] for BACE and [0.65] for BBBP).
B. After Training From Scratch: The middle panels illustrate the embeddings obtained after the MoCL-GAT self-supervised pre-training phase, but before any task-specific fine-tuning. Although the SSL objectives do not explicitly optimize for downstream class separation, some emergent structure begins to appear, reflected in a modest increase in cluster purity (to [0.55] and [0.72]). This suggests the model is capturing general chemical features, but the representations remain substantially task-agnostic.
C. After Fine-tuning (Transfer Learning): The rightmost panels display the embeddings after the pre-trained MoCL-GAT model has been fine-tuned on the labelled data for the specific classification task. A clear improvement in class separability is observed with the blue and orange points forming more distinct clusters. This visual evidence is strongly supported by a significant jump in the purity scores to [0.71] and [0.75]. This demonstrates that fine-tuning effectively adapts the general-purpose representations into task-specific, discriminative embeddings.

Collectively, these visualizations highlight the progression of representation learning in MoCL-GAT. The self-supervised pre-training phase establishes a chemically meaningful embedding space, which is then refined through supervised finetuning to enhance class separability and improve predictive performance on downstream tasks.

### 3.3 Effectiveness of Pre-training

To explicitly quantify the benefits of the self-supervised pre-training stage, we conducted an ablation study comparing the performance of the fine-tuned MoCL-GAT model against an identical GAT architecture trained entirely from scratch on each downstream task. The results, summarized in **Figure *3***, demonstrate that pre-training provides a consistent and significant performance enhancement across all evaluated classification and regression benchmarks. This confirms that the representations learned from large unlabeled datasets are highly transferable and provide a strong inductive bias for supervised learning. For classification tasks, the improvements were particularly pronounced on the BACE dataset, where the AUROC score increased substantially from 0.734 to 0.821, and on BBBP, which rose from 0.898 to 0.928 after fine-tuning.

**Figure 3.**
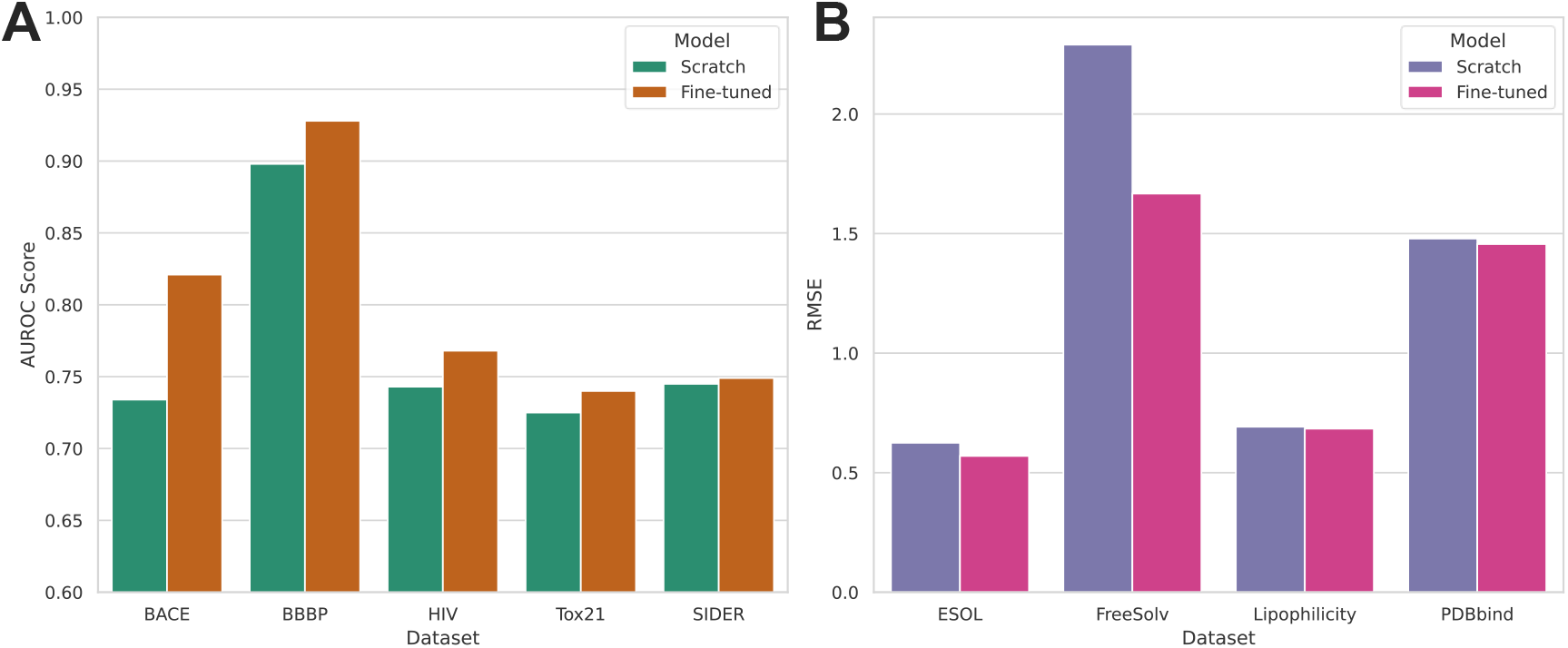
Impact of self-supervised pre-training on downstream task performance. The figure compares MoCL-GAT models fine-tuned after pre-training against identical models trained from scratch. **A)** AUROC scores for classification tasks, <u>where higher is better</u>. **B)** RMSE scores for regression tasks, <u>where lower is better</u>. In all evaluated cases, pre-training and fine-tuning consistently outperforms training from scratch, demonstrating the significant and widespread benefit of the learned representations.

For regression, the most dramatic gain was observed for the FreeSolv dataset, where pre-training lowered the RMSE from 2.290 to 1.667. Even on tasks where the margin was smaller, such as SIDER (AUROC 0.745 to 0.749) or Lipophilicity (RMSE 0.692 to 0.684), the fine-tuned model consistently outperformed the scratch model. These results provide strong quantitative evidence that MoCL-GAT’s dual self-supervised pre-training strategy effectively captures fundamental chemical knowledge, creating a robust foundation that is highly valuable for data-efficient learning on diverse downstream supervised tasks.

Beyond improvements in final predictive performance, pre-training also confers significant advantages in terms of model convergence speed and training stability. Figure S1 illustrates these benefits by plotting the training dynamics on the ESOL dataset. The fine-tuned MoCL-GAT model (blue lines) begins training from a substantially lower initial loss and RMSE across all metrics, a direct result of its knowledgeable initialization from pre-training. It then converges to its optimal performance in significantly fewer epochs compared to the model trained from scratch (red lines), which requires a much longer period to learn basic chemical features from its random initialization. Furthermore, the convergence of the finetuned model is notably more stable, as evidenced by the lower volatility in its validation and test RMSE curves. This demonstrates that transfer learning not only leads to a better final model but also enables a more efficient and reliable training process.

In summary, the experimental results provide compelling evidence for the effectiveness of the MoCL-GAT framework. Quantitative comparisons demonstrate state-of-the-art or highly competitive performance on diverse molecular benchmarks. Qualitative visualizations illustrate the emergence of structured and task-adaptable embeddings. Finally, the finetuning MoCL-GAT confirms the significant advantages conferred by self-supervised pre-training strategy, validating its role in enhancing model generalization and transferability.

## 4 Discussion

In this study, we introduced MoCL-GAT, a novel self-supervised learning framework designed to generate informative and transferable molecular representations using Graph Attention Networks. Our experimental results demonstrate that MoCL-GAT achieves state-of-the-art or highly competitive performance across a diverse range of molecular property prediction and DTI-related benchmark tasks (**Error! Reference source not found**.). The core strength of MoCL-GAT lies in its dual self-supervised learning strategy, which synergistically integrates information from both local atomic environments and global molecular characteristics.

The local contrastive learning component, based on K-hop subgraph sampling and attribute masking, encourages the GAT encoder to learn representations that are robust to minor structural perturbations and focused on the essential chemical features within an atom’s neighborhood. This is particularly valuable for tasks where specific chemical motifs or interaction patterns are critical, such as bioactivity prediction (e.g., BACE, HIV) or side effect profiling (SIDER). The strong performance on SIDER supports the effectiveness of this local objective.

Simultaneously, the global descriptor prediction task compels the model to encode holistic molecular properties, such as overall size, polarity (TPSA), lipophilicity (MolLogP), flexibility (NumRotatableBonds), and druglikeness (QED, SA Score), into the learned representations. This global perspective is essential for predicting pharmacokinetic properties like blood-brain barrier penetration (BBBP), solubility (ESOL), or lipophilicity, where overall molecular features often play a dominant role. The outstanding performance of MoCL-GAT on the BBBP, ESOL, and FreeSolv benchmarks highlights the value of incorporating this global learning signal. We hypothesize that the complementary nature of the local and global objectives enables MoCL-GAT to learn and generate richer and more balanced representations than methods focusing predominantly on a single scale.

The use of GATs as the core encoder further enhances the framework’s effectiveness. The attention mechanism allows the model to adaptively focus on the most relevant parts of the molecular graph for both the local contrastive and global prediction tasks, potentially leading to more expressive embeddings compared to non-attentive GNN architectures. The significant performance gap observed between the fine-tuned MoCL-GAT model and the same architecture trained from scratch (Section 3.3) underscores the value of the self-supervised pre-training phase. The learned representations capture fundamental chemical knowledge that transfers effectively to downstream tasks, reducing the reliance on large-labeled datasets. This transferability is visually supported by the t-SNE analysis (**Figure *2***), which illustrates the evolution from unstructured initial embeddings to structured, task-discriminative representations after pre-training and fine-tuning.

Compared to existing methods (**Error! Reference source not found**.), MoCL-GAT especially effective in predicting pharmacokinetic properties (BBBP, ESOL, FreeSolv) and side effects (SIDER). While specialized models may achieve top scores on individual benchmarks (e.g., MolCLR on BACE, KPGT on Tox21/Lipophilicity, ChemProp on PDBbind), MoCL-GAT offers robust and well-rounded performance across a broader spectrum of tasks, reflecting the versatility of its dual learning approach.

Despite these promising results, this study has several limitations. First, MoCL-GAT currently operates solely on 2D molecular graph representations derived from SMILES, omitting explicit 3D conformational information, which is often critical for modeling molecular interactions, particularly in DTI tasks. Second, the use of a fixed set of 78 global descriptors, while curated, may limit generality; exploring alternative descriptor sets or learning global properties implicitly could yield further improvements. Third, the computational cost of pre-training on nearly 2 million compounds, though performed offline, can be substantial. Lastly, while performance on established benchmarks is strong, validation on prospective, real-world drug discovery datasets or experimental validation of key predictions would further strengthen the claims of practical utility. Additionally, biases inherent in large chemical databases like ChEMBL may influence the learned representations. These biases can be traced to systematic artifacts in the historical drug-discovery process, which include: (1) Scaffold Bias, where chemical scaffolds from historically successful drug campaigns (e.g., kinase inhibitors) are heavily over-represented, potentially leading the model to perform better on common structures than on truly novel ones; (2) Activity Bias, where the database is skewed towards bioactive, “drug-like” compounds, with a significant under-representation of inactive molecules that are crucial for learning effective decision boundaries; and (3) Research Interest Bias, where data is concentrated in well-funded therapeutic areas (e.g., oncology) over others (e.g., rare diseases). A model pre-trained on such data may mistakenly learn these biases, which could affect its generalization performance when applied to chemical spaces or biological targets outside of these well-explored domains.

Future work could address these limitations by incorporating 3D conformational data, potentially through equivariant GNN architectures or complementary pre-training tasks, and by exploring adaptive or learnable global property objectives beyond fixed descriptor sets. Investigating model compression techniques or more efficient pre-training strategies could improve scalability. Finally, applying MoCL-GAT embeddings to challenging downstream applications such as *de novo* molecule generation, reaction prediction, or large-scale virtual screening, coupled with prospective experimental validation, will be crucial for demonstrating its real-world impact.

## 5 Conclusion

In this paper, we presented MoCL-GAT, a novel self-supervised learning framework that leverages Graph Attention Networks to learn powerful molecular representations from unlabeled data. MoCL-GAT uniquely integrates a local contrastive learning objective based on augmented K-hop subgraphs with a global task predicting relevant molecular descriptors. This dual approach enables MoCL-GAT to effectively capture chemical information at multiple scales. Our extensive experiments demonstrate that pre-training with MoCL-GAT generates highly transferable representations, achieving state-of-the-art or strongly competitive performance when fine-tuned on a wide range of benchmark datasets for molecular property prediction and DTI-related tasks. These results highlight the advantages of our dual self-supervised approach and highlight the value of pre-training in addressing the challenge of limited labeled data. MoCL-GAT represents a significant advancement in molecular representation learning, offering a robust and data-efficient tool with strong potential to accelerate computational drug discovery and cheminformatics research. Future work incorporating 3D structural information and prospective validation holds promise for further enhancing its capabilities and practical, real-world impact.

